# eSPRESSO: a spatial self-organizing-map clustering method for single-cell transcriptomes of various tissue structures using graph-based networks

**DOI:** 10.1101/2020.12.31.424948

**Authors:** Tomoya Mori, Toshiro Takase, Kuan-Chun Lan, Junko Yamane, Cantas Alev, Kenji Osafune, Jun Yamashisa, Wataru Fujibuchi

## Abstract

Animal cells are spatially organized as tissues and cellular gene expression data contain such information that governs body structure and morphogenesis during developmental process. Although several computational tissue reconstruction methods using transcriptomic data have been proposed, those methods are insufficient with regard to arranging cells in their correct positions in tissues or organs unless spatial information is explicitly provided. Here, we propose eSPRESSO, a powerful *in silico* three-dimensional (3D) tissue reconstruction method using stochastic self-organizing map (stochastic-SOM) clustering, together with optimization of gene set by Markov chain Monte Carlo (MCMC) framework, to estimate the spatial domain structure of cells in any topology of tissues or organs from only their transcriptome profiles. We confirmed the performance of eSPRESSO by mouse embryo, embryonic heart, adult cortical layers, and human pancreas organoid with high reproducibility (success rate = 72.5–100%), while discovering morphologically important spatial discriminator genes (SDGs). Furthermore, we applied eSPRESSO to analysis of human adult heart diseases by virtual gene knockouts, and revealed candidate mechanisms of deformation of heart structure. The eSPRESSO may provide novel methods to analyze the mechanisms of 3D structure formation and morphology-based disease mechanisms.

## INTRODUCTION

Analysis of disease mechanisms based on high-throughput single-cell RNA-sequencing is becoming prominent and fundamental technique in cell biology^1–3^. Particularly, methods for high-throughput, spatially resolved single-cell RNA-seq have been developed and attracting attentions as novel analytics for disease mechanisms. Single-molecule fluorescence *in-situ* hybridization (FISH)^4^ has been widely used to quantitate transcript numbers at single-cell resolution, often within the context of a diseased tissues of interest. More recently, highly-multiplexed methods such as seqFISH^5^ or merFISH^6^ are reported to detect transcript dynamics of target cells in 3D location. More directly, high-resolved, 2D-mapped primer-based RNA-sequencing of tissues fixed on a glass plate has been also developed for mapping of transcript abundance, utilized to draw spatial cellular location map after reconstructing 3D image by multiple 2D-maps^7^. However, these methods are still in the early stage and requires further research and development in terms of practical cost and conveniences.

Alternatively, several computational methods to reconstruct 3D tissues by estimating the spatial positions of individual cells in tissues with gene expression data obtained by single-cell RNA-seq have been reported.^8–14^ These methods may be roughly divided into two types: the landmark approach and the *ab initio* approach. The landmark approach estimates the 3D position of each cell based on gene expression profiles while using the spatial information of marker genes obtained by *in situ* hybridization.^8–10^ Conversely, the *ab initio* approach assigns each cell to 3D space according to the principal component score calculated from gene expression profiles without using such spatial reference data.^11–14^ Although current principal component analysis (PCA)-based methods are simply used for 3D visualization, an *ab initio* approach that does not depend on the spatial information of marker genes obtained by *in situ* hybridization is promising for 3D reconstruction.

Previously, we reported a novel 3D reconstruction method using stochastic-SOM clustering, or SPRESSO (SPatial REconstruction by Stochastic-SOM)^15^, which features gene selections based on Gene Ontology (GO).^16,17^ The method yielded high success rates of 3D reconstructions of mouse gastrula embryo and demonstrated a remarkable ability to find SDGs that contribute to differentiation and tissue morphogenesis. This method, however, was preliminary and simply projected four domains of mouse gastrula to a cubic structure and thus unapplicable to more complicated structures of organs such as heart or pancreas.

In this paper, we introduced graph-based SOM clustering to reconstruct any topology of domains or tissues as long as they can be drawn as some kind of connection graphs or network diagrams. The concept of graph-based SOM was already reported in 1989 by T. Kohonen and colleagues^18^ who introduced a Kohonen map or network in 1982^19^, which is one of the artificial neural networks and computationally convenient abstraction building on biological models of neuronal systems^20^ and morphogenesis models by Alan Turing^21^. So far, many useful topology structures other than square grids such as toroidal grids that have opposite edges are connected^22^ have been introduced. However, graph-network representation is the most simple but comprehensive for describing relationships tissue domains. Thus, we applied graph-SOM to various types of mouse and human organs that show specific topologies of tissue domain relationships. We also successfully visualized the reconstructed relationships by uniform manifold approximation and projection (UMAP)^23^ to confirm the original structure.

## RESULTS

All the dataset used in this paper is summarized in Table 1 and the detailed results are shown in Supplementary Figures and Tables.

**Table 1.** Datasets and performances of 3D structure reconstructions (score = accuracy + ARI)

| Data Name | Reference | #Domain | #Cells | #Initial genes | #SDGs | Max Accuracy | ARI |
| --- | --- | --- | --- | --- | --- | --- | --- |
| Mouse embryo (E7.0) | Peng, Guangdun, et al. "Spatial transcriptome for the molecular annotation of lineage fates and cell identity in mid-gastrula mouse embryo." *Developmental cell* 36.6 (2016): 681-697. | 4 | 41(sections) | 185 | 18 | 100.0% | 1 |
| Mouse embryo (E5.5-7.5) | Peng, Guangdun, et al. "Molecular architecture of lineage allocation and tissue organization in early mouse embryo." *Nature* 572.7770 (2019): 528-532. | 7 | 213 (sections) | 87 | 17 | 86.3% | 0.79731 |
| Mouse brain (ALM) | Tasic, Bosiljka, et al. "Shared and distinct transcriptomic cell types across neocortical areas." *Nature* 563.7729 (2018): 72-78. | 3 | 3809 | 106 | 7 | 100.0% | 1 |
| Mouse brain (VISp) |  | 4 | 7049 | 122 | 19 | 100.0% | 1 |
| Mouse heart (E7.75) | de Soysa, T. Yvanka, et al. "Single-cell analysis of cardiogenesis reveals basis for organ-level developmental defects." *Nature* 572.7767 (2019): 120-124. | 4 | 1259 | 51 | 9 | 100.0% | 1 |
| Mouse heart (E8.25) |  | 7 | 3331 | 132 | 22 | 97.5% | 0.93082 |
| Mouse heart (E9.25) |  | 7 | 3911 | 93 | 18 | 96.8% | 0.92909 |
| Mouse liver | Halpern, Keren Bahar, et al. "Single-cell spatial reconstruction reveals global division of labour in the mammalian liver." *Nature* 542.7641 (2017): 352-356. | 9 | 1415 | 97 | 18 | 84.6% | 0.48085 |
| Human pancreas (S4) | Veres, Adrian, et al. "Charting cellular identity during human in vitro β-cell differentiation." *Nature* 569.7756 (2019): 368-373. | 5 | 5273 | 107 | 21 | 95.9% | 0.86908 |
| Human pancreas (S5) |  | 5 | 3925 | 51 | 21 | 97.6% | 0.94836 |
| Human pancreas (S6) |  | 5 | 5051 | 56 | 18 | 95.5% | 0.853 |
| Human heart (4.5-5PCW) | Asp, Michaela, et al. "A spatiotemporal organ-wide gene expression and cell atlas of the developing human heart." *Cell* 179.7 (2019): 1647-1660. | 8 | 238 (spots) | 57 | 37 | 76.2% | 0.5314551 |
| Human heart (6.5PCW) |  | 8 | 1515 (spots) | 108 | 15 | 70.6% | 0.3526241 |
| Human heart (9PCW) |  | 8 | 1358 (spots) | 101 | 24 | 72.5% | 0.3513529 |

### Spatial clustering of gene expression data by graph-based SOM with gene set optimization

Inspired by Kohonen’s self-organizing-map learning theory^19^, here we extended our previously developed method, SPRESSO^15^, to graph-based networks, which is theoretically applicable to cell-to-cell relationships of any types of topological structures of tissues or organs. The basic algorithm of spatial clustering of cells is a combinatorial optimization to find best gene sets to reproduce known topological network structures of learning objects, or gene vectors of cells. Since each node on the topological network may contain multiple such gene vector objects, given an original topological network, we can calculate the reproducibility of the structure by counting correct and incorrect network edges, either connected or unconnected, after learning process. We also used the Markov-Chain-Monte-Carlo (MCMC) to optimize the best gene sets to give a maximum reproducibility of a topology of structure. We have also tuned the SOM learning process by introducing stochastically learning (i.e., stochastic-SOM^15^) that allows learning efficiency in later phase where the extent of learning ability usually decreases monotonously. We refer to this method as eSPRESSO (enhanced-SPRESSO) in this paper.

Here we first tested the ability of eSPRESSO with mouse E5.5-7.5 gastrula embryo, whose topological structures are too complicated to be handled and reproduced by our previous SPRESSO, by generating graph structure of 7 domains (Ect1, Ect2, Ect3, PS, MA, MP, E1-E2-E3) with domain-to-domain contact relationships (Fig.1a). We optimized the gene set to maximize the reproducibility of the original graph structure by 1,000 steps of MCMC calculations. To enhance the efficient optimization and stable reproducibility, we performed parallel tempering^24^ based on 8 parallel MCMC processes. The resulted clustering is evaluated with the original structure by simply counting the existing edges between two gene vector objects, or cells. We used gene expression data from 213 sections and the reproduced accuracy was 86.3% after the MCMC optimization and the domain contact map (Fig.1b) of samples for original (lower triangle) and reconstructed (upper triangle) structures (Fig.1c) are shown. A heatmap of gene expression data for optimized SDGs (Fig.1d) is also shown. The final clustering using SDGs is drawn by UMAP (Fig.1e).

**Figure 1.**
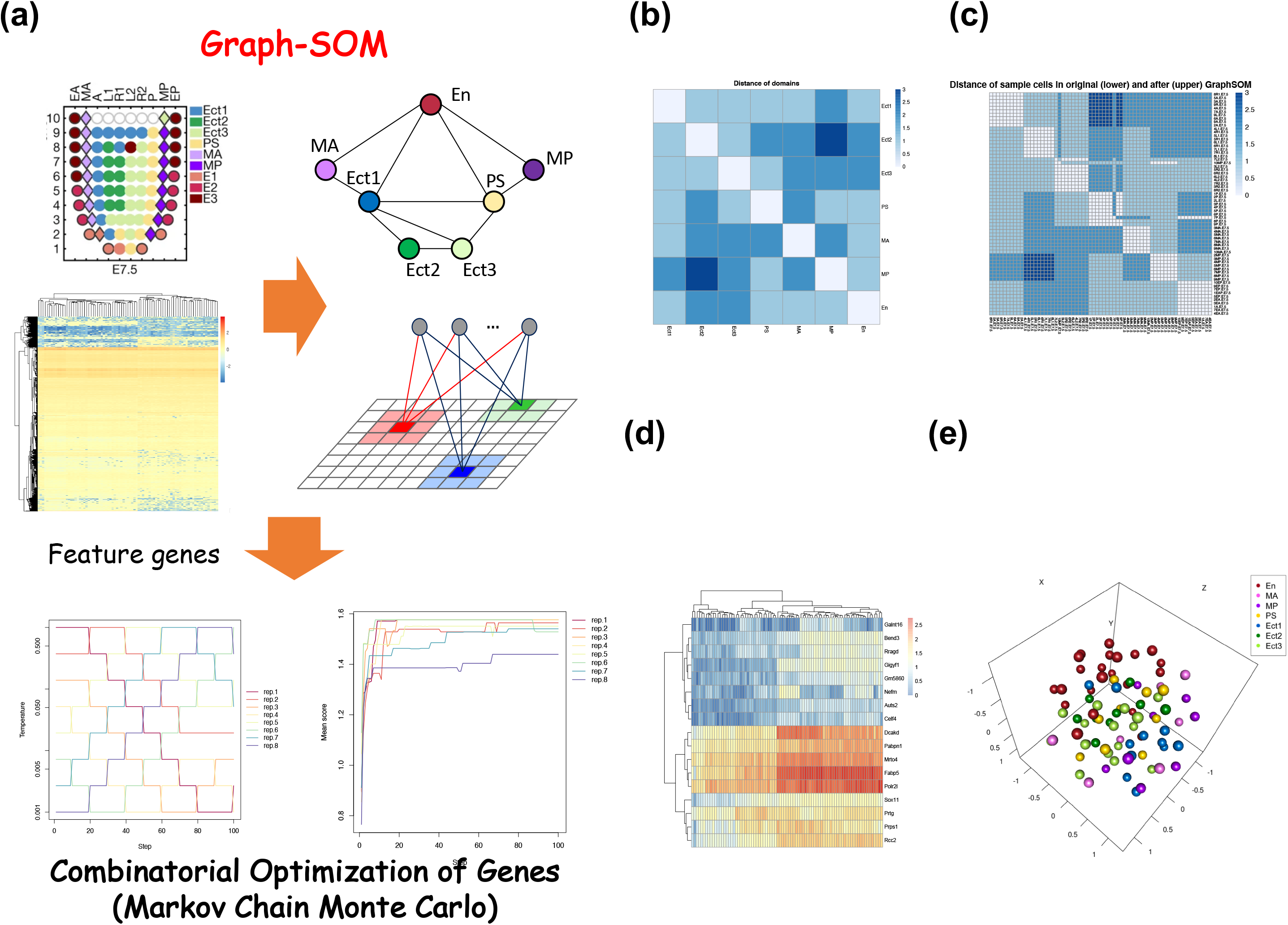
Overview of the 3D reconstruction method of mouse embryo using stochastic-SOMclustering with graph-network topology.

### Circular and hierarchical structure reconstruction by eSPRESSO

To evaluate the performance of eSPRESSO, we applied the method to circular and hierarchical structures of tissues. The development of mouse heart initiates with an expansion of a blood vessel, forming complex structure consisting of arteries and veins in later phase. We used scRNA-seq data from mouse embryonic heart structure (E9.25) which represents pseudo-circular structure of seven domains, of which five are projecting from the anterior and posterior second heart fields (AHF and pSHF) (Fig.2a). When we down-sampled 10 cells for each domain from the initial 3911 cells and optimized the gene set by stochastic-SOM with MCMC to reproduce the original structure, we successfully reconstructed the topology of cellular locations with extremely high accuracy of 96.8% (Fig.2b). The number of final SDGs is 18 which contains many known important genes such as *Hand2*, *Gata4*, etc.^25^, in mouse heart development. The visual projection by UMAP clustering of all 3331 cells using the SDGs show a circular representation of tissue domains which is quite consistent of original structure (Fig.2a).

**Figure 2.**
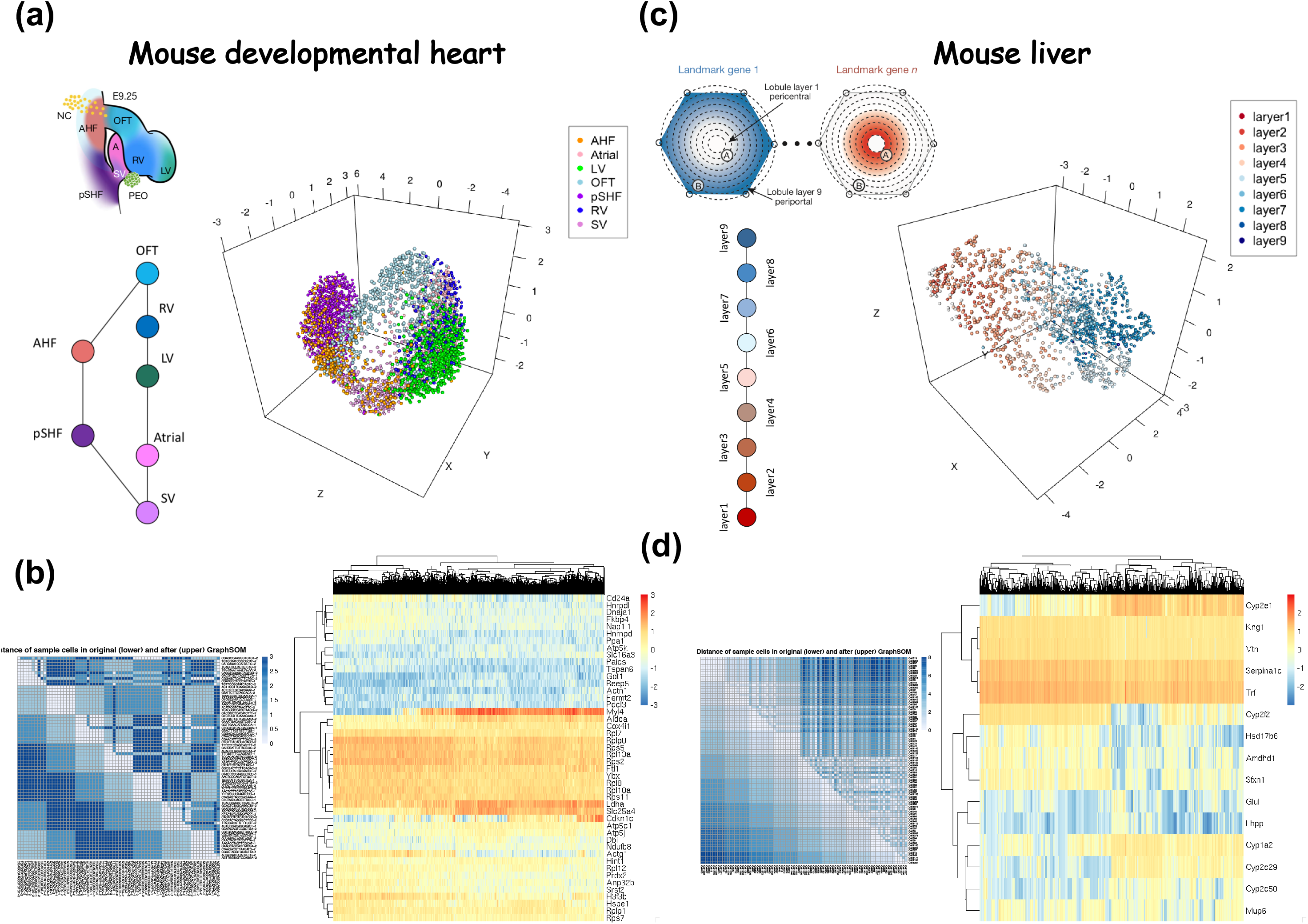
Examples of 3D reconstructions: a,b) mouse developmental heart and c,d) mouse liver.

Then we further tried to reconstruct mouse liver structure where a nine layers of tissues from a concentric circle are observed^10^ (Fig.2c). The eSPRESSO correctly reconstructed the relationships of 90 cells, which are randomly selected by down-sampling 10 cells from each of 9 domains, with 84.6% accuracy. The resulted 18 SDGs represent many known liver-specific genes such as cytochrome P450 (*Cyp1a2* and *Cyp2f2*) and hepatic nuclear factor 4 alpha (*Hnf4a*)^27^ (Fig.2d). The visual projection by UMAP clustering of all 1415 cells using the 18 SDGs indicate that there are clear layer structure observed although it does not reproduce the circular structure due to the limitation of UMAP clustering that are difficult to generate circular layer topology (Fig.2c).

### Developmental analysis of human pancreas by eSPRESSO

One of the powerful characteristics of eSPRESSO may be demonstrated by analysis of organs during developmental process. We used scRNA-seq data from human developmental pancreas from stage 4, 5 and 6 data^28^, and reconstructed the structure. The accuracies are high for all the stages such as 96-100%. The numbers of resulted SDGs are 9, 22, and 18, containing pancreas-specific genes such as insulin (*INS*) and insulin gene enhancer protein ISL-1 (*ISL1*)^29^ (Supplementary Table1). Interestingly, although we did not input information of endocrine and non-endocrine separation, the UMAP plots of all the thousands of cells using the SDGs indicate that developmental events such as alpha-cell differentiation in Langerhans islets and exocrine cell detachment from endocrine cells are observed (Fig.3a-c).

**Figure 3.**
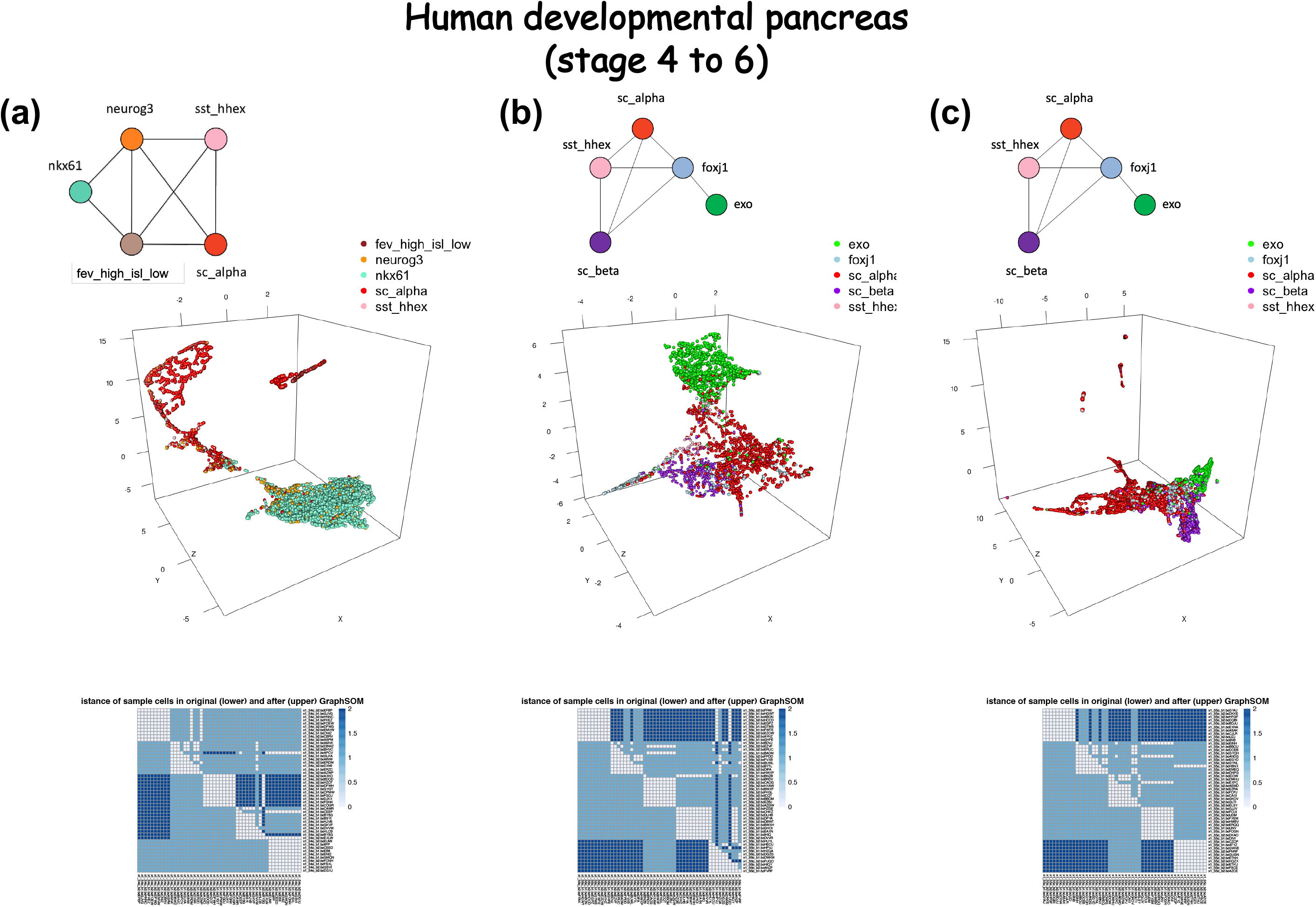
Example of 3D reconstructions during pancreas development.

In the stage 4 to 6 reconstructions, the original scRNA-seq data is focusing on endocrine cells; therefore, we obtained endocrine-related genes during stages. GSEA analysis also represent that the developmental changes of SDGs, indicating that more polarization between endo- and exocrine cells seemed to proceed.

### Application to virtual human gene knockout experiment

One of the most useful usages of eSPRESSO may be to perform virtual *in-silico* human gene knockout experiments. As an example of organs, we used scRNA-seq data from human developmental heart^30^. We first successfully reconstructed the 8 domain structure with accuracy of 76.2% for postconceptional weeks (PCWs) 4.5-5 human embryonic heart (Fig.4a). We obtained ## SDGs as the optimized result (Supplementary Table1). Then we performed single gene deletions among the SDGs to see if any structural changes are observed (Fig.4b). Among the 37 SDGs, most effective genes that dramatically decreases accuracies of reconstructions are found (Supplementary Table2). The structural change for gene X deletion is shown in Fig.4c. The gene X is the orthologousgene in mouse that is homozygous lethal. The decreases of accuracies are also shown (Supplementary Table2). All other structural changes are shown in Supplementary Figure 1. We also performed two gene deletions (Supplementary Table3).

**Figure 4.**
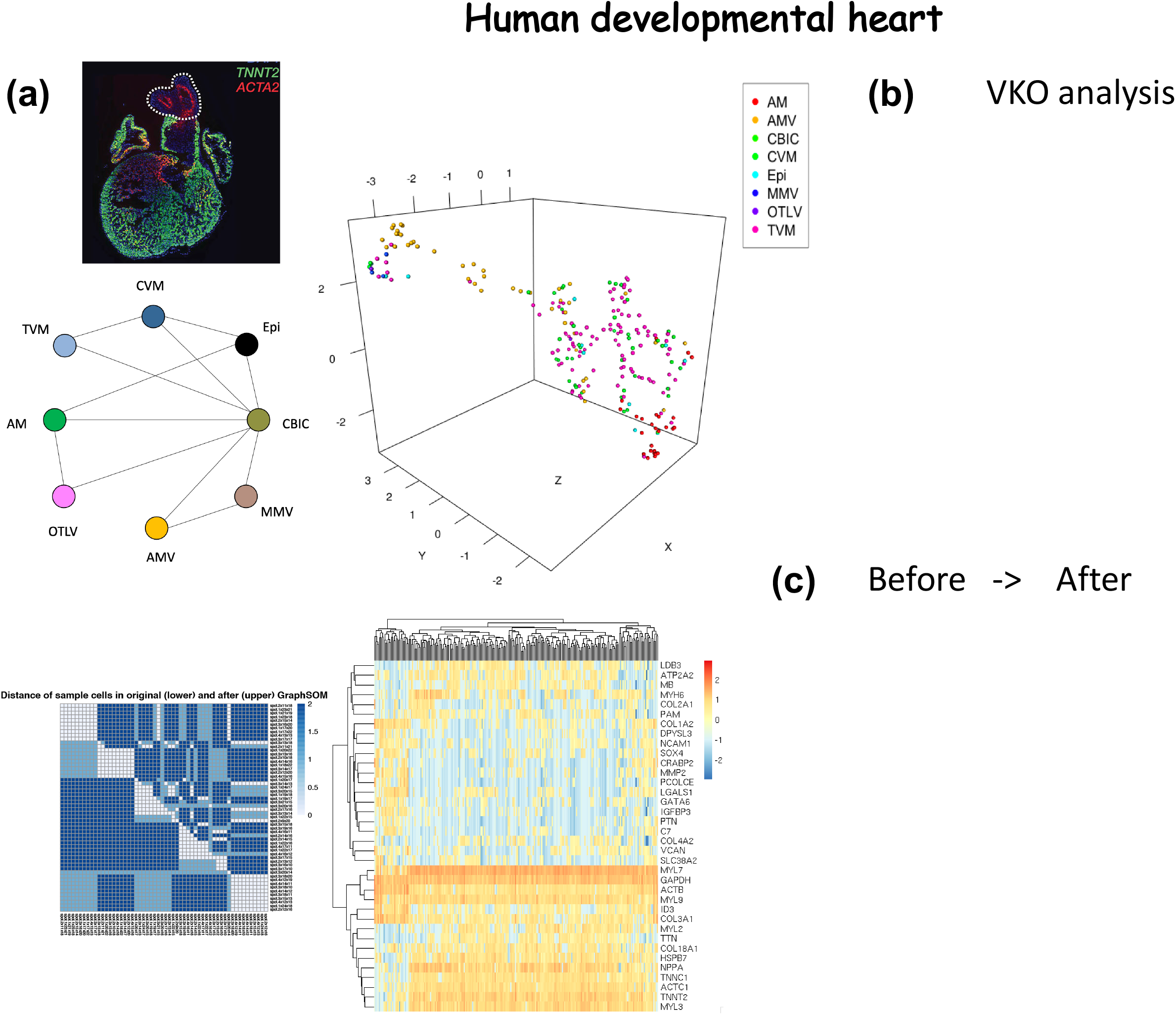
Example of virtual knockout analysis by eSPRESSO in human developmental heart.

We also performed similar analyses for scRNA-seq data from PCW 6.5 and 9 human hearts. The most effective genes that decreases accuracies of reconstructions are shown in Supplementary Table 4.

## Discussion

In this paper, we developed a computational approach that combines the SOM clustering under the topological constraints and gene set optimization by MCMC. The reproducibility of tissue topology is sufficiently high and this method may innovatively add more information on the existing clustering method such as helping to find spatially distributed discriminator genes (SDGs) and inference of developmental architecture of tissue domains. It is also useful to investigate the collapses of spatial clustering by *in-silico* virtual gene knockouts although it is not a real knock-out experiments.

Recently, alternative direct experimental approaches such as highly-multiplexed FISH or direct spatial transcriptome have been proposed. However, the primal purpose of these methods is to reconstruct cellular locations with gene expression data. Although they can contribute to detect differentially expressed genes among distinct tissue domains, it does not explicitly consider the tissue topologies and thus, it may lack to detect spatially contributing genes in terms of neighboring tissue networks. Our method may be able to complement such a neighboring tissue information and it is also applicable to the results of such data to detect globally expressed genes that contributes to domain networks in spatial distributing manner

Our methods may have benefits in analysis of organ development. Many genes show homozygous lethality in development. Although many evidences based on animal studies indicate that there are candidates of those lethal genes in human, it is not possible to reproduce lethality in human system due to ethical reasons. The human organoids derived from iPS or ES cells may provide an alternative approach to reproduce gene knockout organs. However, abnormalities are often stochastic and affected by other internal or external factors and it sometimes requires a large number of repeat experiments to reproduce the phenomena^31^. Contrastingly, our *in-silico* virtual knockout method does not require any cumbersome protocols and may be able to reproduce the same results in many times.

The SOM learning is a renowned method in the field of clustering, first proposed by Kohonen and colleagues^19^. They have already introduced graph-network topology for SOM clustering but there existed not many examples to solve in their days^18^. The original SOM learning decreases learning regions and efficiency monotonously, which causes clustering problems in the later phase and we found that efficient scheduling for learning was necessary for practical application to the current single-cell transcriptome datasets. Thus, we introduced the stochastic-SOM learning which has a characteristic similar to that of Gibbs-sampling although the sampling probability is totally equal. This improvement dramatically enhanced the clustering ability and thus enabled to implement this study.

Finally, we state that our current algorithms and analysis pipeline describe here is still primitive and many modifications are necessary. The extraction of initial gene candidates are solely dependent on Boruta package that detects feature genes by random forest generation and may miss important genes that overlaps categories in the distant branches or leaves. The parallel tempering MCMC is also a suboptimal approach, although it enhances the combinatorial optimization to allow quick convergence to the global optimum, which sometimes fail to find the best gene set to reproduce the topology by SOM clustering. We expect that more efficient and fine-grained algorithms to find global optimum in decent time may replace modules in our pipeline in future.

## Methods

**Methods and any associated references are available in the online version of the paper**.

**Note: Any Supplementary Information and Source Data files are available in the online version of the paper**.

## Code and data availability

The proposed methods including the feature gene selection, the 3D reconstruction using stochastic-SOM clustering, and visualization, are implemented in R and Python and are available at https://github.com/tmorikuicr/espresso. Also all the gene expression data used in this study is included together.

## Acknowledgements

The authors deeply appreciate Dr. Peter Karagiannis for kindly reviewing the manuscript. This work was partially supported by the Core Center for iPS Cell Research, Research Center Network for Realization of Regenerative Medicine, Japan Agency for Medical Research and Development (16bm0104001h0004), and the iPS Cell Research Fund.

## Author contributions

W.F. conceptualized and designed the study. T.M. implemented the software. T.M., K.L. and T.T. performed the computational experiments. J.Yamane., C.A., K.O., and J.Yamashita. performed the data interpretation from a biological point of view. T.M. and W.F. wrote the manuscript. All authors have read and approved the final manuscript.

