## Supplementary File for "eSPRESSO: a spatial self-organizing-map clustering method for single-cell transcriptomes of various tissue structures using graph-based networks"

**Supplementary Information**

**Data collection and preprocessing**

**Data collection and preprocessing**

To confirm the performance of eSPRESSO, 14 datasets of human and mouse transcriptome were collected from seven papers [1-7]. Each dataset was used as log10-transformed expression profile and the genes whose expression values are greater than 1.0 in at least two samples and whose standard deviation across all samples are greater than 0.05 were extracted. Adjacent matrices of spatial domains were construct from original papers and biological knowledge by the authors of this paper.

**Mm embryo (E7.0)**

Peng *et al*. collected transcriptome profiles on embryo section by laser-microdissection [1]. We downloaded and used E1 dataset from [GSE65924](https://www.ncbi.nlm.nih.gov/geo/query/acc.cgi?acc=GSE65924), which is one of the triplicates of single embryos. The dataset contains 41 sections (~20 cells per sample) with the four spatial domains (d1:Anterior, d2: lateral-distal, d3: lateral-proximal, and d4: posterior). The expression values were saved as FPKM.

**Mm embryo (E7.5)**

Peng *et al*. collected transcriptome profiles on embryo section of various developmental stages by laser-microdissection [2]. We downloaded reference samples of E7.5 from [GSE120936](https://www.ncbi.nlm.nih.gov/geo/query/acc.cgi?acc=GSE120963). The dataset contains 83 sections (20-40 cells per samples) with the nine spatial domains (Ect1-3: ectoderm, PS: primitive streak, MA: anterior mesoderm, MP: posterior mesoderm, E1-3: endoderm). To simplify the domain structure, we integrated E1-3 into En as a single domain. The expression values were saved as FPKM.

**Mm brain (ALM and VISp)**

Tasic *et al*. collected transcriptome profiles on two regions of adult mouse cortex: anterior lateral motor cortex (ALM) and primary visual cortex (VISp) [3]. We downloaded exon counts datasets of ALM and VISp from [GSE115746](https://www.ncbi.nlm.nih.gov/geo/query/acc.cgi?acc=GSE115746), respectively. For both datasets, we extracted 3,809 and 7,049 single cells of L2/3, L4, L5, L6, and L6b clusters, respectively, where the L6b cells were merged to the L6 cluster. The ALM consists of three spatial domains of L2/3, L5, and L6, whereas the VISp consists of the four spatial domains of L2/3-L6. The expression values were saved as raw count data, and they were transformed to CPM values when analyzed.

**Mm heart (E7.75, E8.25, and E9.25)**

Yvanka de Soysa *et al*. collected transcriptome profiles on mouse heat of three developmental stages: E7.75, E8.25, and E9.25 [4]. We downloaded source data of all developmental stages from supplementary files of the original paper. In addition, we requested expression profiles of sinus venosus (SV) and Atria of E9.25 from the authors of the original paper to integrate to the downloadable E9.25 data. For E7.75 data, the anterior heart field (AHF), the left ventricle (LV), Atrial, and the posterior second heart field (pSHF) of wild type were extracted and the total number of single-cells was 1,259 while 3,331 and 3,911 single-cells of AHF, the SHF-derived outflow tract (OFT), the right ventricle (RV), LV, Atrial, SV, and pSHF were extracted from E8.25 and E9.25 data. The expression values were saved as log-transformed UMI counts.

**Mm liver**

Halpern *et al*. collected transcriptome profiles on mouse liver and estimated their lobule coordinates by a panel of zonated landmark genes [5]. The authors provided the single-cell gene expression profile and a posterior probability matrix showing the probabilities of being the original layer for each single-cell, and they can be downloaded as supplementary files of the original paper. Here, in order to simplify the input data, we determined the layer that gives the maximum probability as the original layer for each single-cell. This dataset consists of 1,415 single-cells with the nine domains (layer1-9). The expression values were saved as raw UMI counts and it was transformed to CPM when analyzed.

**Hs pancreas (S4, S5, and S6)**

Veres *et al*. collected transcriptome profiles on human pancreas of four differentiated stages [6]. We downloaded the gene expression profile of stages 4-6 (S4-6) from [GSE114412](https://www.ncbi.nlm.nih.gov/geo/query/acc.cgi?acc=GSE114412), where profiles of replications were removed. For S4, the expression profiles are consist of 5,273 single-cells with the five domains: NKX6-1^+^ progenitors (nkx61), NEUROG3^+^ progenitors (neurog3), SC-$\alpha$ (sc_alpha), SST^+^HHEX^+^ (sst_hhex), and FEV^high^ISL^low^ (fev_high_isl_low). For S5 and S6, the original datasets consist of six domains: SC-$\beta$ (sc_beta), SC-$\alpha$ (sc-alpha), SC-EC (sc-ec), CHGA^+^FOXJ1^+^ (foxj1), SST^+^HHEX^+^, and Non-endocrine (exo). However, it is difficult for S5 and S6 data to uniquely determine their adjacency matrices since the connection between SC-EC and the other domains are ambiguous. Therefore, here we removed SC-EC domain and transformed S5 and S6 datasets to five domains data. The expression values of these datasets were saved as UMI counts and they were transformed to CPM when analyzed.

**Hs heart (4.5-5 PCW, 6.5 PCW, and 9 PCW)**

Asp *et al*. collected transcriptome profiles on human heart at three developmental stages in the first trimester: 4.5-5, 6.5, and 9 post-conception weeks (PCW) [7]. Each expression profile and corresponding metadata are able to downloaded from a data repository (<https://www.spatialresearch.org>). The spatial transcriptome data of 4.5-5 PCW, 6.5 PCW, and 9PCW were consists of 238, 1,515, and 1,358 spots of tissue sections. The 4.5-5 PCW and 6.5 PCW data consist of the eight domains: compact ventricular myocardium (CVM), trabecular ventricular myocardium (TVM), atrial myocardium (AM), outflow tract and large vessels (OTLV), atrioventricular mesenchyme and valves (AMV), mediastinal mesenchyme and vessels (MMV), cavities with blood and immune cells (CBIC), and epicardium (Epi) while the 9 PCW data consists of the seven domains excepting CBIC.

**Stochastic self-organizing map (stochastic-SOM) clustering**

Self-organizing map (SOM) is an unsupervised clustering method proposed by Kohonen [8]. In general, SOM projects input high-dimensional data onto a limited number of output classes or units, so that the different units with similar centroid vectors are placed close to each other in a mapping layer, which is usually given in a two-dimensional (2D) plane. Let $\boldsymbol{X}=\left( \boldsymbol{x}_{1},\boldsymbol{x}_{2},\ldots,\boldsymbol{x}_{n} \right)$ is a set of input samples with the $p$-dimensional vectors, i.e., $\boldsymbol{x}_{j}=\left( x_{j1},x_{j2},\ldots,x_{jp} \right)$ $\left( j=1, 2,\ldots, n \right)$. The mapping layer is consist of $k$ units, and their centroid vectors $\boldsymbol{m}_{i}=\left( m_{i1}, m_{i2},\ldots,m_{ip} \right)$ $\left( i=1, 2, \ldots, k \right)$ is randomly initialized and assigned to each unit. The similarity between input sample $j$ and all units $i$ is defined by the Euclidean distance. First, the SOM algorithm finds the unit $c$ with the highest similarity according to the following equation as the best matching unit (BMU).

$$c=\arg\min_{i\in\left\{ 1,\ldots,k \right\}} \left\{ \left\| \boldsymbol{x}_{j}-\boldsymbol{m}_{i}\left( t \right) \right\| \right\},$$

where $\left\| \cdot\right\|$ denotes the Euclidean distance, or norm of a vector and $t$ is time step. The centroid vector $\boldsymbol{m}_{i}\left( t \right)$ of all units of the mapping layer at time $t$ is updated by the following equations.

$$\boldsymbol{m}_{i}\left( t+1 \right)=\boldsymbol{m}_{i}\left( t \right)+h_{ci}\left( t \right)\left( \boldsymbol{x}_{j}-\boldsymbol{m}_{i}\left( t \right) \right),$$

$$h_{ci}\left( t \right)=\alpha\left( t \right)\exp\left( -\frac{\left\| \boldsymbol{r}_{c}-\boldsymbol{r}_{i} \right\|^{2}}{2\sigma^{2}\left( t \right)} \right),$$

where $h_{ci}\left( t \right)$ is a neighborhood function which determines how much $\boldsymbol{m}_{i}\left( t \right)$ receives the learning influence of $\boldsymbol{x}_{j}$ when it is updated. $\alpha\left( t \right)$ and $\sigma\left( t \right)$ are the learning rate parameter and a function defining the radius of the neighboring region, respectively. In addition, $\boldsymbol{r}_{c}$ and $\boldsymbol{r}_{i}$ are position vectors in the mapping layer of unit $c$ and $i$. The SOM algorithm repeats updates of $\boldsymbol{m}_{i}$ until the learning step $t$ reaches $T$ which is given as a parameter for all input samples $j$.

In the general SOM clustering, its result is affected by the order in which the samples are input. To remove the effect, the batch-learning SOM was also proposed in [8]. In the batch-learning SOM, $\boldsymbol{m}_{i}\left( t \right)$ is updated only after all samples are given by the following equations.

$$c_{j}\left( t \right)=\arg\min_{i\in\left\{ 1,\ldots,k \right\}} \left\{ \left\| \boldsymbol{x}_{j}-\boldsymbol{m}_{i}\left( t \right) \right\| \right\},$$

$$\boldsymbol{m}_{i}\left( t+1 \right)=\frac{\sum_{j=1}^{n} h_{c_{j}\left( t \right)i}\left( t \right)\boldsymbol{x}_{j}}{\sum_{j=1}^{n} h_{c_{j}\left( t \right)i}\left( t \right)}.$$

The general SOM learning often converges to local minima in early steps if the number of units in the mapping layer is extremely small. In order to increase the possibility of escaping from the local minima and reaching the global maxima, a stochastic-SOM that introduces random variable into the neighborhood function has been proposed, which make the learning converge gradually [9]. The neighborhood function of the stochastic-SOM is

$$h_{ci}\left( t \right)=\alpha\left( t \right)\exp\left( -\frac{\mathrm{rnd}\left[ 0.5, 1 \right)\cdot\left\| \boldsymbol{r}_{c}-\boldsymbol{r}_{i} \right\|^{2}}{2\sigma^{2}\left( t \right)} \right),$$

where $\mathrm{rnd}[0.5, 1)$ is a function generates uniform random value at least 0.5 and less than 1.0. In this paper, the learning rate $\alpha\left( t \right)$ is set to be 1.

**Graph-based stochastic-SOM clustering**

A graph $G=(V,E)$ is a pair of a finite non-empty set $V$ and finite set $E\subseteq V\times V$. The elements $u,v\in V$ of the graph $G$ are called vertices, and the elements $e=\left\{ u,v \right\}\in E$ are called edges. The set of vertices and edges of a graph $G$ are denoted as $V(G)$ and $E(G)$, and the number of them are denoted as $|V(G)|$ and $|E(G)|$, respectively. For all edges $\{u,v\}\in E\left( G \right)$, if $(u,v)=(v,u)$ then the graph $G$ is called an undirected graph. Hereinafter, a graph $G$ is an undirected graph unless specified. A path on $G$ is a non-empty graph $P(G)=(V,E)$, where $V=\left\{ v_{i},v_{i+1},\ldots,v_{j} \right\}$ and $E=\left\{ \left\{ v_{i},v_{i+1} \right\},\left\{ v_{i+1},v_{i+2} \right\},\ldots,\left\{ v_{j-1},v_{j} \right\} \right\}$, and all $v_{k}$ are distinct. The distance $d_{G}\left( u,v \right)$ between two vertices $u$ and $v$ on $G$ is given by the length of the shortest path between $u$ and $v$. A graph $G$ is often represented by a square matrix $A\left( G \right)=\left[ a_{ij} \right] \left( i,j=1,2,\ldots,\left| V\left( G \right) \right| \right)$ that shows the adjacency between vertices, which is called an adjacency matrix, where $a_{ij}=1$ if $\left\{ v_{i},v_{j} \right\}\in E\left( G \right),$ otherwise $a_{ij}=0$ for the vertices $v_{i}$ and $v_{j}$ corresponding to $i$ and $j$, respectively.

Here, in order to improve the performance of the stochastic-SOM, we newly propose a graph-based SOM (graph-SOM) clustering. In the graph-SOM, the mapping layer is given by a graph $G$ represented by the adjacency matrix $A\left( G \right)$. Although, in the general SOM, the distance between units $i$ and $j$ in the mapping layer is computed by the Euclidean distance $\left\| \boldsymbol{r}_{i}-\boldsymbol{r}_{j} \right\|$ between the corresponding position vectors $\boldsymbol{r}_{i}$ and $\boldsymbol{r}_{j}$, each unit corresponding to a vertex on $G$ and the distance between the units is given by distance $d_{G}\left( v_{i},v_{j} \right)$ between the vertices $v_{i}$ and $v_{j}$ on $G$ in the graph-SOM. Therefore, the neighborhood function $h_{ci}\left( t \right)$ at time $t$ of the stochastic graph-SOM is given by the following equation.

$$h_{ci}\left( t \right)=\alpha\left( t \right)\exp\left( -\frac{\mathrm{rnd}\left[ 0.5, 1 \right)\cdot d_{G}\left( v_{c},v_{i} \right)^{2}}{2\sigma^{2}\left( t \right)} \right).$$

**Optimization of gene set by Markov chain Monte Carlo (MCMC) framework**

In order to get the spatial discriminator genes (SDGs), eSPRESSO employs Random Forest-based feature gene selection method Boruta [10] and replica exchange Markov chain Monte Carlo-based gene set optimization.

**Feature gene selection by Boruta**

Kursa and Rudnicki proposed Boruta method for a Random Forest based algorithm for feature selection [10]. The one of the properties of Boruta is to classify features into three classes: *confirmed*, *tentative*, and *rejected* rather than order them. In eSPRESSO clustering, we selected around 100 *confirmed* genes as features by increasing the value of a parameter `maxRuns` in the Boruta package of a programming language R.

**Replica exchange MCMC optimization**

After getting feature genes by Boruta, eSPRESSO searches for the optimum combination of the feature genes by replica exchange MCMC framework and then output SDGs. The replica exchange MCMC is an extended algorithm of MCMC for improving the sampling efficiency [11]. In a general simulated annealing (SA) algorithm [12], which is one of the optimization algorithms based on MCMC sampling, there is only one temperature parameter that determines whether to adopt or reject the newly obtained sample, and the probability of being adopted is relatively high even for samples with the large energy difference when the temperature is high. However, it is difficult to get out of the local minima at low temperature. As a result, the probability of being rejected increases and sampling efficiency decreases. In the replica exchange MCMC, multiple systems called replicas with different parameters are simulated at the same time, and the state of the replicas are exchanged between different temperatures according to the following exchange probability;

$$p=\min_{} \left( 1,\exp\left( \left( E_{i}-E_{j} \right)\left( \frac{1}{T_{i}}-\frac{1}{T_{j}} \right) \right) \right),$$

where $E_{k}$ and $T_{k}$ are energy and temperature of replica $k$. By updating the variables of each replica at their respective temperatures and moving on the temperature axis at the same time, its long-term behavior can be regarded as a random walk.

In eSPRESSO, sampling in each replica is done by the following MCMC sampling algorithm.

(Step 1) Let $G_{b}$ be the feature gene set obtained by Boruta.
(Step 2) Select $n$ genes from $G_{b}$ at uniformly random and let$G’$ be the set of the $n$ genes.
(Step 3) Repeat the following procedures $T$times.
i. Generate candidate gene sets $\mathcal{g=}\left\{ G_{cand}^{1},G_{cand}^{2},\ldots,G_{cand}^{N} \right\}$ by adding a gene $g_{i}\in G_{b}\setminus G'$ to $G’$ for all $g_{i}$ or deleting a gene $g_{j}\in G^{'}$ from $G’$ for all $g_{j}$.
ii. Remove gene sets $G_{cand}^{i}$, which are already sampled.
iii. If $\mathcal{g}$ is an empty set, replace a gene $g_{j}\in G'$ with the other genes $g_{k}\in G_{b}-\{g_{j}\}$ and add them to $\mathcal{g}$.
iv. For all $G_{cand}^{i}$, execute the stochastic graph-SOM clustering and compute the scores of the clustering results (see the section **Evaluation of graph-SOM clustering results** for more details).
v. Define a selection probability $p_{i}$ for $G_{cand}^{i}$ as

$$p_{i}=\frac{\left( \exp\left( z_{i} \right) \right)^{c}}{\sum_{i=1}^{N} \left( \exp\left( z_{i} \right) \right)^{c}},$$

where $z_{i}=\frac{s_{i}-\mu}{\sigma}$, and $\mu$ and $\sigma$ are the mean value and the standard deviation of the scores $s_{i}$ of $G_{cand}^{i}$. $c$ is a constant parameter and $c=\sqrt{\left| \mathcal{g} \right|/2}$ is employed in the computational experiments in this paper.

vi. Determine whether to adopt or reject of $G_{cand}^{x}$ according to the adoption probability $p_{SA}$ of general simulated annealing (SA) strategy.

$$p_{SA}=\left\{ \begin{aligned} 1 &(\Delta f\leq0) \\ \exp\left( \frac{-\Delta f}{T_{t}} \right) &(\Delta f>0) \end{aligned} \right.$$

Note that eSPRESSO handles the maximization problem for the score $s$, so that the difference $\Delta f$ is defined as $\Delta f=-\left( s_{x}-s \right)$, where $s_{x}$ and $s$ are the scores of $G_{cand}^{x}$ and $G$, respectively.

vii. Update $G^{'}$ by $G_{cand}^{x}$ if adopted, and update also $G$ by $G_{cand}^{x}$ if the score of $G_{cand}^{x}$ is larger than that of $G$.
(Step 4) Output $G$ as the optimized gene set.

**Evaluation of graph-SOM clustering results**

The clustering result of the stochastic graph-SOM is evaluated by two criteria: prediction accuracy and adjusted Rand index (ARI).

**Prediction accuracy**

For a pair of cell samples $c_{i}$ and $c_{j}$, let $d_{i}$ and $d_{j}$ be the true domains to which they belongs, and let $\hat{d}_{i}$ and $\hat{d}_{j}$ be the domains to which they are estimated to belong by the stochastic graph-SOM. Here, assuming that $s_{xy}$ is an element of the adjacency matrix $A(G)$ corresponding to the input graph $G$, the prediction score $s_{ij}$ for a pair $\left\{ c_{i},c_{j} \right\}$ is given by the following equation

$$s_{ij}=\left\{ \begin{aligned} 1 &(a_{d_{i}d_{j}}=a_{\hat{d}_{i}\hat{d}_{j}}) \\ 0 &(a_{d_{i}d_{j}}\neq a_{\hat{d}_{i}\hat{d}_{j}}) \end{aligned} \right..$$

Therefore, the prediction accuracy $s$ for all cell sample pairs is defined by the following equation

$$s=\frac{\sum_{i,j}^{\left( \begin{matrix} n \\ 2 \end{matrix} \right)} s_{ij}}{\left( \begin{matrix} n \\ 2 \end{matrix} \right)},$$

where *n* is the total number of cell samples.

**Adjusted Rand index**

The adjusted Rand index (ARI) measures the similarity between two clustering results [13]. Let $\mathcal{X=}\left\{ \mathcal{X}_{1},\mathcal{X}_{2},\ldots,\mathcal{X}_{m} \right\}$ and $\mathcal{Y=}\left\{ \mathcal{Y}_{1},\mathcal{Y}_{2},\ldots,\mathcal{Y}_{m} \right\}$ be families of sets of cell samples, where $m$ is the number of domains. The overlap of cell samples between $\mathcal{X}_{i}$ and $\mathcal{Y}_{j}$ is denoted by $n_{ij} (=\left| \mathcal{X}_{i}\cap\mathcal{Y}_{j} \right|)$. Thus, the number of cell samples belonging to $\mathcal{X}_{i}$ (resp. $\mathcal{Y}_{j}$) can be represented using $n_{ij}$ as $a_{i}=\sum_{j=1}^{m} n_{ij}$ (resp. $b_{i}=\sum_{i=1}^{m} n_{ij}$). Therefore, the ARI can be defined as the following equation:

$$ARI\left( \mathcal{X,Y} \right)=\frac{\sum_{i,j} \left( \begin{matrix} n_{ij} \\ 2 \end{matrix} \right)-\left[ \sum_{i} \left( \begin{matrix} a_{i} \\ 2 \end{matrix} \right)\sum_{j} \left( \begin{matrix} b_{j} \\ 2 \end{matrix} \right) \right]/\left( \begin{matrix} n \\ 2 \end{matrix} \right)}{\frac{1}{2}\left[ \sum_{i} \left( \begin{matrix} a_{i} \\ 2 \end{matrix} \right)+\sum_{j} \left( \begin{matrix} b_{j} \\ 2 \end{matrix} \right) \right]-\left[ \sum_{i} \left( \begin{matrix} a_{i} \\ 2 \end{matrix} \right)\sum_{j} \left( \begin{matrix} b_{j} \\ 2 \end{matrix} \right) \right]/\left( \begin{matrix} n \\ 2 \end{matrix} \right)},$$

where $n$ is the total number of cell samples. Here, assuming that $\mathcal{X}$ and $\mathcal{Y}$ are the true family of domains and the family of domains estimated by the stochastic graph-SOM, respectively, the similarity between the true domain classification and estimated domain classification can be computed by ARI.
